# Building regulatory landscapes: enhancer recruits cohesin to create contact domains, engage CTCF sites and activate distant genes

**DOI:** 10.1101/2021.10.05.463209

**Authors:** Niels J. Rinzema, Konstantinos Sofiadis, Sjoerd J. D. Tjalsma, Marjon J.A.M. Verstegen, Yuva Oz, Christian Valdes-Quezada, Anna-Karina Felder, Teodora Filipovska, Stefan van der Elst, Zaria de Andrade dos Ramos, Ruiqi Han, Peter H.L. Krijger, Wouter de Laat

**Affiliations:** Oncode Institute, Hubrecht Institute-KNAW and University Medical Center Utrecht, 3584 CT Utrecht, The Netherlands

## Abstract

Developmental gene expression is often controlled by distal tissue-specific enhancers. Enhancer action is restricted to topological chromatin domains, typically formed by cohesin-mediated loop extrusion between CTCF-associated boundaries. To better understand how individual regulatory DNA elements form topological domains and control expression, we used a bottom-up approach, building active regulatory landscapes of different sizes in inactive chromatin. We demonstrate that transcriptional output and protection against gene silencing reduces with increased enhancer distance, but that enhancer contact frequencies alone do not dictate transcription activity. The enhancer recruits cohesin to stimulate the formation of local chromatin contact domains and activate flanking CTCF sites for engagement in chromatin looping. Small contact domains can support strong and stable expression of distant genes. The enhancer requires transcription factors and mediator to activate genes over all distance ranges, but relies on cohesin exclusively for the activation of distant genes. Our work supports a model that assigns two functions to enhancers: its classic role to stimulate transcription initiation and elongation from target gene promoters and a role to recruit cohesin for the creation of contact domains, the engagement of flanking CTCF sites in chromatin looping, and the activation of distal target genes.

## INTRODUCTION

Tissue-specific and developmentally restricted expression of genes is often controlled by enhancers, regulatory DNA elements that can act over distance to activate gene expression. In the mammalian genome, enhancers may be located near their target gene, but they can also be separated over hundreds of kilobases. Chromatin looping is thought to enable distal enhancers to approach and activate target genes. Indeed, individually studied developmental genes were found to form preferred contacts with distal enhancers for their activation^1^, as measured by chromosome conformation capture (3C) methods^2,3^. Genome-wide, a general correlation between enhancer-promoter (E-P) contact frequencies and transcriptional activity was uncovered by these methods^4-7^. Also, non-coding disease-associated genetic variants, identified by genome-wide association studies (GWAS), could be successfully linked to target genes when considering their 3C-based chromatin contacts^8^. As an enhancer was found to activate expression when it was forced to loop to a distant gene, E-P looping seems a requirement for, more than a consequence of, long-range gene activation^9^.

Cohesin is responsible for chromatin loop formation in interphase chromosomes. *In vitro*, the ring-shaped cohesin complex can extrude DNA to form loops^10,11^, exactly as it was predicted to create chromatin loops *in vivo*^12,13^. DNA-bound CTCF is the most dominant factor to bind and halt the DNA extruding cohesin complex on chromatin and to anchor DNA loops in mammalian cells^14-18^. CTCF-anchored loops are the most readily detectable loops in genome-wide Hi-C contact maps^16^, highlighting their frequent recurrence across cells. They span sub-megabase domains, also known as topologically associating domains (TADs), containing sequences that preferentially self-interact, and they form boundaries that hamper contacts between neighboring domains.

Without cohesin, both CTCF loops and contact domains dissolve in Hi-C chromatin contact maps, but overall steady-state gene expression levels seemed remarkably stable^19,20^. This raised doubts about whether chromatin looping was really necessary for enhancers to control gene expression over distance, doubts that were further fueled by live cell microscopy studies that sometimes^21^ but not always^22,23^ found enhancers in closer proximity to the genes that they activated. Yet, other studies did observe that cohesin depletion caused loss of promoter-enhancer contacts^24,25^ and reduced enhancer-dependent expression^24-27^ of many, but not all genes, suggesting that both cohesin-dependent and -independent mechanisms exist for long-range gene regulation^25,28^. Acute depletion of WAPL, the factor that releases cohesin from chromatin, caused repositioning of cohesin from tissue-specific enhancers to CTCF boundaries, disrupted enhancer-promoter looping and down-regulated expression of tissue-specific genes controlled by such enhancers^24^. This suggested that these enhancers can serve as cohesin entry sites and that active cohesin loading is required for their productive interaction with target genes^24^. Support for this model came from findings that NIPBL, the cohesin loading factor, is preferentially associated with enhancers^29,30^ and appeared required for long-range gene regulation^20^. Looping between CTCF sites, which depends on cohesin, was found to correlate with the presence of active promoters and enhancers in the intervening chromatin^31,32^, further suggesting that active regulatory DNA elements can act as entry sites for the loop extruding cohesin machinery.

CTCF boundaries insulate chromatin domains not only physically but also functionally, as they can obstruct enhancers to activate genes in neighboring domains. Consequently, cognate enhancers of a given gene are typically found in the same contact domain^15,33-36^. Within a contact domain, a proximal gene in principle has an advantage over a distal gene for activation by a shared enhancer^36-38^. Promoter mutations that interfere with enhancer contacts can re-direct the enhancer to contact and activate more distal genes^39,40^.

Collectively, this suggests an intricate interplay between enhancers, CTCF sites, and cohesin to form contact domains and control the expression of distant genes. To experimentally investigate this further, we took a bottom-up approach and built a large series of different regulatory landscapes in an inactive chromatin environment. In this chromatin setting, we find that enhancers recruit cohesin to stimulate the building of self-interacting domains and to engage flanking CTCF sites in chromatin looping. Enhancers simultaneously serve to boost and stabilize the transcriptional activity of distant genes, which is more effective at shorter linear distances and in smaller domains. Irrespective of the enhancer-promoter distance, the enhancer relies on cognate transcription factors and mediator for transcription regulatory activity. In contrast, cohesin is exclusively required for long-range transcription activation, not for the enhancer to activate a proximal gene promoter. Our work, therefore, shows that a developmental enhancer can recruit the cohesin loop extrusion machinery to promote longer-range chromatin contacts, build contact domains and enable long-range gene activation. Since the same enhancer-promoter pair relies on cohesin for gene activation strongly when separated over large distances (>100 kb), mildly when apart over intermediate distances (47kb) and not at all in proximal (<11kb) configurations, our data shows that linear distance importantly determines whether an enhancer requires cohesin for ‘long-range’ gene regulation.

## RESULTS

### Increasing enhancer-promoter distances reduce transcriptional activity and stability

To investigate the role of regulatory DNA sequences in building transcriptional regulatory landscapes and forming topological domains, we experimentally searched for a suitable chromatin environment. This, we reasoned, had to be a transcriptional neutral or repressive chromosomal segment. As regulatory DNA sequences, we used the 6.5 kb human β-globin micro-LCR^41^ (μLCR), a prototype of a strong tissue-specific enhancer, and a β-globin gene (*HBG1*) promoter-driven GFP-reporter as its target gene (**Fig. 1a**). We randomly integrated them as a single construct in the genome of erythroleukemia K562 cells and selected high GFP-expressing clones. We then used Cre recombinase to remove the μLCR in order to find clones whose high reporter gene expression strictly relied on the μLCR. These clones, we reasoned, had integrated the transgene in a transcriptional non-supportive chromatin environment. The integration sites were mapped and with help of publicly available Hi-C and ChIP-seq datasets we selected an integration site on chromosome 18 (Chr18: 19609009) inside a relatively large (nearly 600kb) and diffuse structural chromatin domain that was covered with repressive H3K27me3 histone marks, had few small H3K27Ac sites and lacked expressed genes. The right boundary of this domain, 500kb away from the integrated reporter gene, showed a striped pattern in Hi-C, indicative of anchored loop extrusion activity^30^. Elsewhere in the locus, a few selected CTCF sites showed cohesin association, implying that this locus was not devoid of natural cohesin association and activity (Fig 1b).

**Fig.1.**
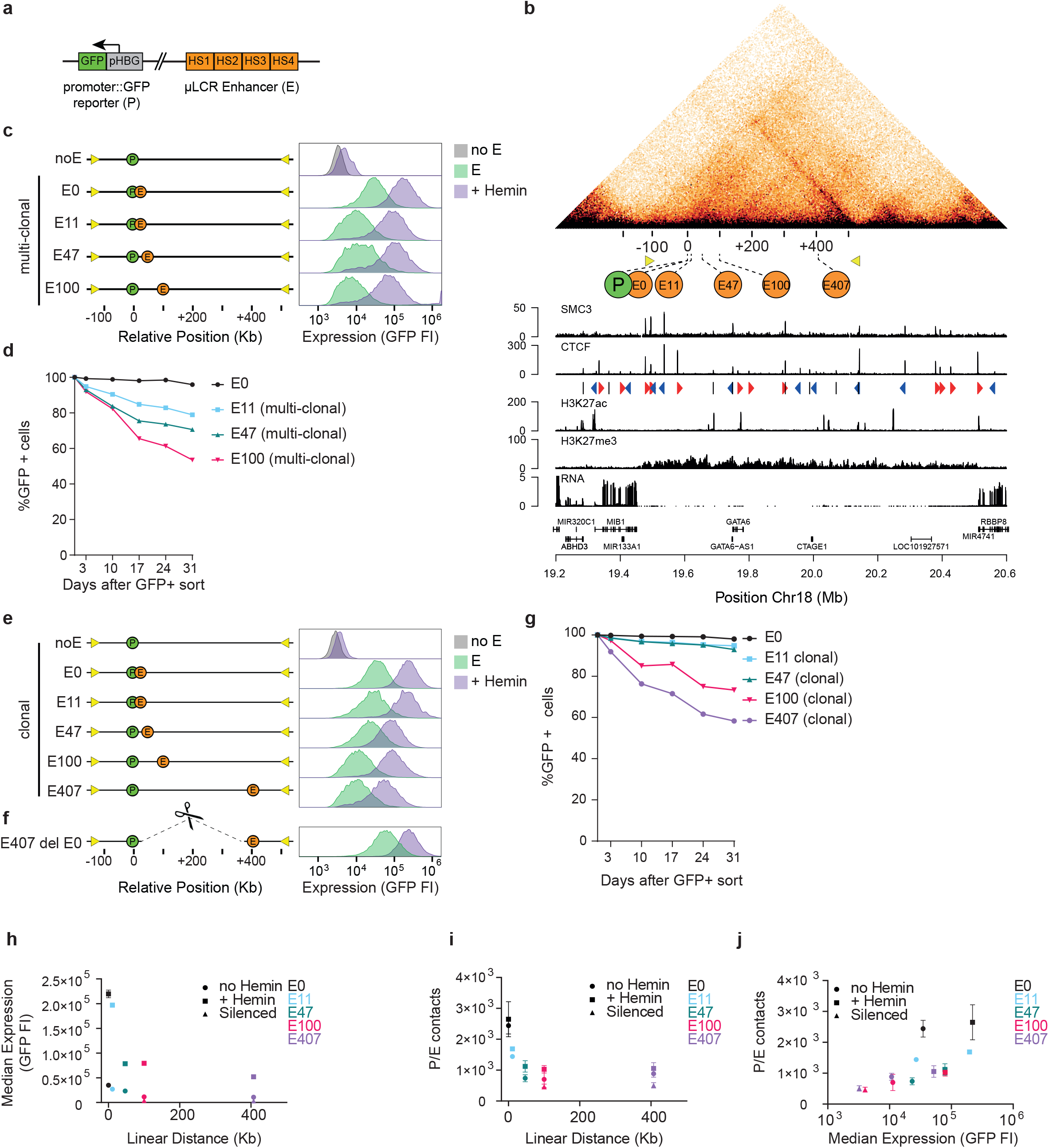
Enhancer-promoter distances and contact frequencies versus gene expression. **a**. Schematic representation of Promoter (P) / Enhancer (E) pair. The reporter gene consists of the human Hemoglobin Subunit Gamma 1 (HBG1) promoter driving GFP expression. The µLCR enhancer consists of four DNase I hypersensitive sites (HS1-4) of the human β-globin locus control region (LCR)^41^. **b**. Genomic context of integrated transgene in K562 cells: Knight-Ruiz (KR) normalized HiC contact map at 5 Kb resolution taken from^16^, with genomic scale (kb) centered on the integrated transgene (P), and with the different enhancer integration sites (E) indicated. For reference, yellow triangles demarcate the ‘left boundary’(see text) and right domain boundary. Tracks below from top to bottom show: ChIP-seq tracks^63^ for SMC3, CTCF (red triangles: forwardly orientated CTCF Binding Sites (CBS), blue triangles: reversely orientated CBSs), H3K27ac, H3K27me3, RNA-seq, and Ref-seq genes. Genomic positions on chromosome 18 are shown below in Mb. **c**. GFP Fluorescence Intensity in arbitrary units (FI (a.u.)) of multi-clonal (bulk) cell populations carrying no enhancer (noE) or the µLCR enhancer at 0 kb, 11 kb, 47 kb or 100 kb (E0, E11, E47, E100, respectively) upstream of the GFP-reporter, without hemin (green) or with hemin (purple) in the culture medium. **d**. Percentage of remaining GFP positive cells in multi-clonal (bulk) cell populations carrying the enhancer at indicated distances, after long-term cell culturing. At each time point, cells were first treated with hemin for two days prior to FACS analysis. **e**. Same as in **c**, but for selected clonal cell lines carrying the µLCR enhancer at indicated distances **f**. GFP Fluorescence Intensity (FI) of an E407 cell line, modified by the deletion of intervening sequences to have the µLCR enhancer placed at 0 kb of the reporter gene. **g**. Same as in **d**, but for selected clonal enhancer cell lines (E-lines) carrying the µLCR enhancer at indicated distances. **h-j**. Median reporter gene expression (GFP fluorescence) plotted against linear E-P distance (**h**), and mean 4C-seq-measured E-P contacts (mean normalized (per 1 million *cis*-reads) 4C-seq signal at enhancer) plotted against E-P linear distance (**i**) or against Median GFP fluorescence (**j**). Cells were either not treated with hemin (circles), treated with hemin (squares) or long term silenced (triangles). Color-code corresponds to that used in **g**. Error bars represent standard deviation (n= two technical replicates/clone).

We then first studied the impact of enhancer-promoter (E-P) distance on transcriptional output. For this, we used CRISPR-Cas9 to re-insert the μLCR at 0, 11, 47 and 100 kb upstream of the reporter gene. Bulk analysis of GFP-expressing cells demonstrated that transcriptional output decreased with increasing E-P distance (**Fig. 1c**). K562 cells can be stimulated by hemin to further differentiate towards the erythroid lineage^42^ and to upregulate LCR-mediated globin gene expression^43^. At all E-P distances, hemin treatment resulted in a roughly five-fold further upregulation of the reporter gene. This showed that the inverse relation between linear E-P distance and transcriptional output was maintained under conditions that stimulated enhancer communication (**Fig. 1c**).

Expansion of GFP positive sorted cells revealed that some cells lost their ability to express GFP over time, even if they were stimulated by hemin. To further investigate this, we again site-specifically inserted the μLCR at the selected distances and bulk sorted cell populations for GFP expression for two consecutive weeks. We then FACS monitored them weekly, each time after a two day hemin induction. When immediately flanking the reporter gene (at 0 kb distance), the enhancer conferred long-term stable expression: nearly all cells (>95%) remained active, even after 5 weeks of culturing. With increasing distance, however, the enhancer correspondingly lost capacity to protect the reporter gene from silencing (∼20% (E11), 30% (E47) and 50% (E100) silenced cells after 5 weeks) (**Fig. 1d and Extended Data Fig. 1a**). The linear distance between the enhancer and promoter in a repressive chromatin environment, therefore, related inversely not only to transcriptional activity, but also to transcriptional stability.

### Enhancer-promoter contact frequencies do not strictly dictate transcriptional activity

Given that chromatin contact frequencies generally decay exponentially with increased chromosomal distance, our data, similar to other work^36^, suggested a relationship between enhancer-mediated gene activation and E-P contact frequencies. We used 4C-seq^44^ to more directly analyze E-P contact frequencies. 4C-seq involves quantifying the competitive ligation events between a genomic site of interest and its spatially most proximal DNA fragments inside each cell nucleus, which then provides a semi-quantitative measure for contact frequencies in a cell population. We first selected for each E-P distance pair a clonal cell line (the enhancer lines, or E-lines) that we genetically confirmed and validated to display representative transcriptional activity and stability (**Fig. 1e, g**). We failed to generate bulk (multi-clonal) populations with the enhancer at 200, 300 or 400 kb (even if we co-integrated a CTCF boundary, see below), but managed to sort out an individual E-line expressing GFP under control of the μLCR at 407 kb. This E407 cell line expressed GFP at lower levels and was also more prone to silencing than clones having a more proximal enhancer (**Fig. 1e, g**), further showing that transcriptional activity and stability decreased with enhancer distance. When we deleted the >400 kb intervening sequence in long-term silenced E407 cells (i.e. cultured for more than 6 weeks, weekly selected for the absence of GFP expression), rare cells with very high GFP expression were identified and clonally expanded (**Fig. 1f**). Genotyping confirmed they carried the deletion that placed the enhancer at 0 kb distance from the reporter gene. This further supported that enhancer distance can impact expression levels and suggested that gene silencing can be reversed when moving the enhancer closer to the gene, as previously observed through forced chromatin looping^9^. Note though that not all cells with the deletion re-expressed the reporter gene at high levels. We applied 4C-seq to the reporter gene to analyze its chromatin contacts in the different E-lines. As a proxy for E-P contact frequencies, we quantified the contacts with the μLCR and expressed it as a percentage of intra-chromosomal 4C contacts. When plotted against the average GFP expression levels, we observed an overall positive but not a linear relationship between E-P contact frequencies and transcriptional activity (**Fig. 1h-j**). To further investigate this relationship, we took advantage of the ability of our system to modulate expression levels at a given enhancer position. First, we performed similar 4C-based measurements after hemin treatment, which induced reporter gene expression in all E-lines roughly 5-fold. This upregulation was generally accompanied by a slight increase in E-P contact frequencies (**Fig. 1h-j, and Extended Data Fig. 2a**). We then investigated E-P contacts in long-term silenced clones (see above) that no longer expressed the reporter gene, even not after hemin induction. Here, E-P contact frequencies were slightly reduced as compared to active cells (**Fig. 1 h-j, and Extended Data Fig. 2b**). Collectively these results support other recent data that the relationship between gene activity and E-P contact frequencies is non-linear and that subtle changes in contact frequencies can lead to large changes in expression^36,45,46^.

### Tissue-specific enhancer forms local self-interacting chromatin domains

We then asked whether the integration of the reporter gene or enhancer had an impact on the topology of the locus. To test this for the reporter gene, we assayed the chromatin contacts of the integrated transgene promoter in cells lacking a co-integrated enhancer and compared these to contacts made by the endogenous sequence immediately flanking its integration site in wild type (WT) cells. Contact profiles were almost identical (**Fig. 2a, and Extended Data Fig. 3a**), which suggested that insertion of the reporter gene itself had little impact on chromatin topology.

**Fig2.**
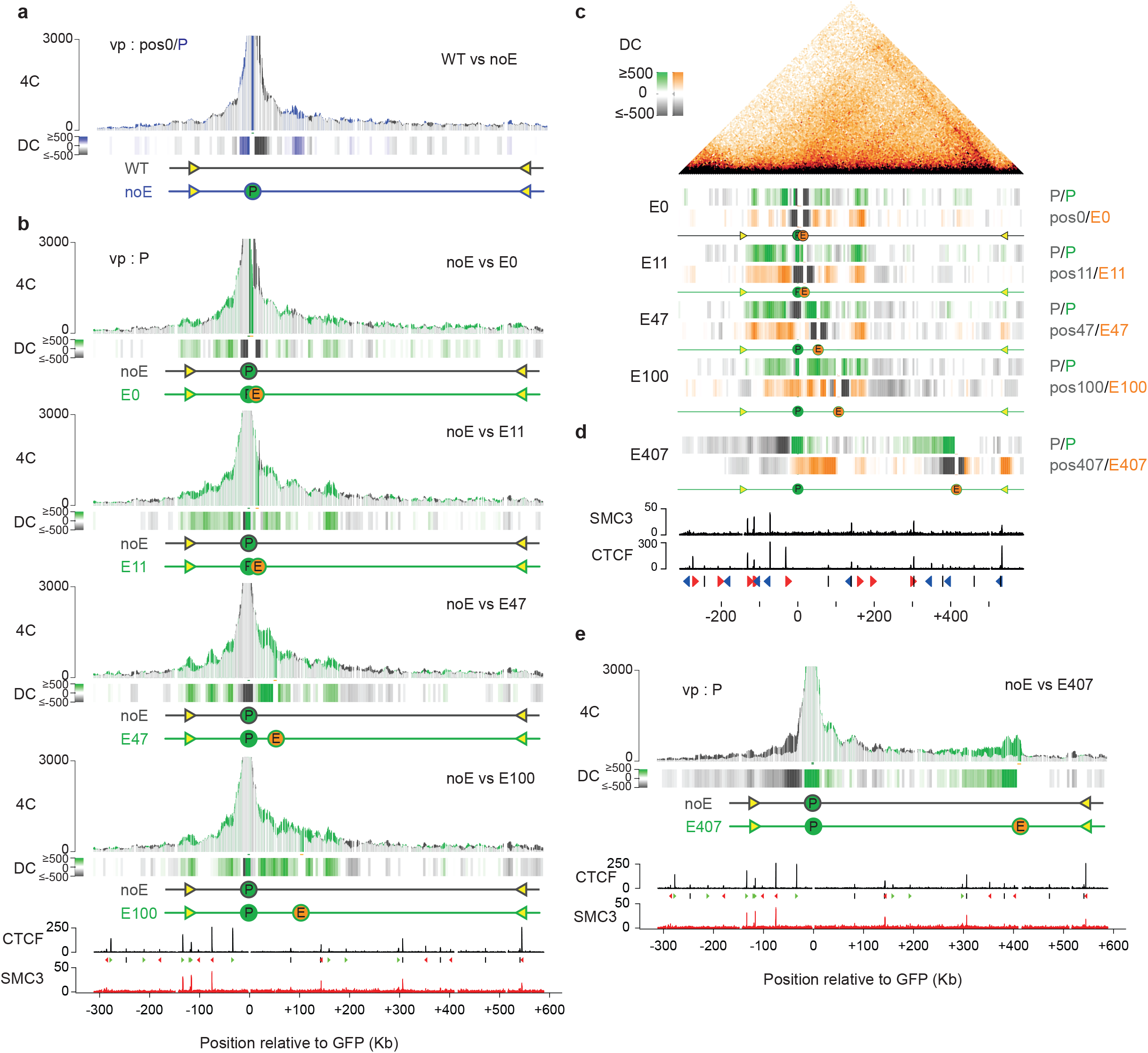
Tissue-specific enhancer forms local self-interacting chromatin domains. **a**. 4C-seq contact profiles plotted as overlays, comparing contacts of the integrated reporter gene promoter (P) in a cell line lacking the enhancer (noE, in blue), versus contacts of the corresponding endogenous genomic position (Pos0) in wildtype (WT, in dark grey) K562 cells. Shared contacts are in light grey. Y-axis: 4C coverage per 1 million *cis*-reads. VP: 4C-seq viewpoint. Track below shows the differential 4C-seq signal (DC: differential contacts), with contacts gained and lost by the gene promoter in blue and grey, respectively. n = two technical replicates/clone. **b**. 4C-seq profile overlays comparing contacts of the integrated promoter (P) in the different E-lines (green profiles), versus those of the integrated promoter in the cell line lacking an integrated enhancer (noE: dark grey profiles). Differential contacts (DC) are plotted below each overlay profile. Bottom: CTCF and SMC3 ChIP-seq tracks. **c**. 4C-seq differential contacts (DC) tracks showing, per E-line (E0, E11, E47 and E100), the gained (green) and lost (gray) contacts of the gene promoter as compared to its contacts in the cell line lacking the enhancer (noE), as well as the gained (orange) and lost (grey) contacts of the enhancer, as compared to its corresponding endogenous chromosomal position in the cell line lacking the enhancer (noE). For reference, the aligned HiC contact map is shown on top. VP: 4C-seq viewpoint. **d**. same as in **c**, but for the E407 cell line. SMC3 and CTCF ChIP-seq signal and CTCF-site orientations are indicated at bottom. **e**. 4C-seq overlay contact profiles showing gene promoter contacts gained (green) and lost (black) upon integration of the enhancer at position E407, as compared to its contacts in a cell line carrying no integrated enhancer (noE).

In contrast, when we plotted the reporter gene’s 4C contact profiles in the E-lines as overlays over its contacts when it had no integrated μLCR in *cis*, it became obvious that the distantly located enhancer stimulated the gene to engage bi-directionally in longer-range contacts. Gene contacts were stimulated not only with the enhancer itself, but also with intervening and surrounding sequences, across a defined genomic interval (**Fig. 2b**). This genomic interval of enhancer-induced gene contacts appeared similar between all lines having the enhancer integrated within 100 kb, and corresponded to the domain that each of these integrated enhancers preferentially contacted themselves (**Fig. 2c, Extended Data Fig. 3b**). A different enhancer-activated contact domain was observed in the E407 line. Here, the ultra-far upstream enhancer exclusively stimulated contacts of the reporter gene with upstream sequences, across 400kb of sequences towards and with the upstream enhancer (**Fig. 2d, e**). The E407 enhancer itself probed this same region, with distal interactions with the region just upstream of the gene, and two prominent more specific contacts, one with an undefined intervening DNA element and the other with the flanking right CTCF-boundary (**Extended Data Fig 3b**). Collectively the data show that an enhancer at different locations can stimulate a gene to engage in different chromatin contacts and induce the formation of different local contact domains.

### Smaller domains support enhancer-induced expression levels and stability

We wished to study the relationship between domain sizes and gene regulation in more detail, and introduced a 3x-hCTCF cassette, with three strong CTCF-binding sites (CBS) selected from the human genome. We successfully integrated the μLCR together with the 3x-hCTCF cassette at 0, 11, 47 and 100 kb from the reporter gene, with the CBSs oriented convergently, facing the enhancer and (distal) reporter gene (we failed to find integrations at 407 kb). We generated bulk GFP-positive cell populations as before and found that at all four integration sites, the presence of flanking CBSs resulted in higher levels of transgene expression, even after hemin induction (**Fig. 3a**). Furthermore, we found that the flanking CBSs also helped the enhancer at all distances to protect the transgene from silencing (**Fig. 3b**). We then selected representative clonal cell lines with the enhancer-3x-hCTCF cassette at 0 kb (EC0) and 100 kb (EC100) (EC-lines). As expected, 4C-seq demonstrated that the ectopic CBSs served as boundaries: they hampered gene promoter and enhancer contacts across the integration sites but stimulated their contacts with downstream sequences to effectively create a smaller and more self-interacting domain containing the enhancer and reporter gene (**Fig. 3c**). Taking advantage of lox sites flanking the μLCR, we removed the enhancer in EC0 and EC100 to create C0 and C100 cell lines, (C-lines), having only ectopic CBS (no enhancer) integrated in *cis* with the reporter gene. The C-lines showed that the ectopic CBS itself had no intrinsic transcription activation capacity, nor did they stimulate the reporter gene to engage in new chromatin contacts, as the enhancer did (**Fig. 3d**). In fact, it was the enhancer that stimulated the integrated CBS to form a stable chromatin loop with a convergent endogenous CTCF site (termed the ‘left boundary site’). This was most notably seen in the EC0 line, but was also appreciable in the EC100 line (**Fig. 3d and Extended Data Fig. 4a**). We then asked whether a 3xCTCF cassette at the gene, instead of the enhancer, had similar impact. We created an E100 cell line (C-0E100) with a 3xCTCF cassette located downstream of and convergent to the gene. In C-0E100, the CBS gave some, but much less support to transcription than in EC100, where the CBS was placed upstream of the enhancer (**Fig. 3e**, compare to **Fig. 3c**). We liked to attribute this to the fact that the CTCF sites downstream of the gene failed to find a proximal looping partner, with the gene and enhancer consequently still acting in a larger and less insulated domain (**Fig. 3e**). However, we also considered the option that a CBS may act differently when flanking a gene or an enhancer. To further investigate this we placed a dsRed reporter gene instead of the enhancer at position 100 kb, in a dual reporter cell line that also had the original GFP reporter gene at 0 kb and the enhancer downstream at respectively 50 kb and 150 kb of each of the genes. We then modified this locus further to also have this regulatory landscape acting with a CBS at 100 kb, flanking the dsRed reporter gene (**Fig. 3f**). Here, the juxtaposition of CBS to the gene now also strongly increased the enhancer-mediated transcriptional output per cell and the ability to express over time. (**Fig. 3g**). Thus, no matter whether it flanked the gene or the enhancer, a CBS introduced at position 100 that placed the E-P pair in a much smaller contact domain, strongly supported long-range enhancer-mediated transcription, while the CBS flanking the gene at position 0 that had little impact on the domain size, did not obviously support transcription. We therefore like to speculate that CTCF sites that create smaller domains enable distal enhancers to confer increased transcriptional activity and stability to genes, possibly because they concentrate loop extrusion activity.

**Fig3.**
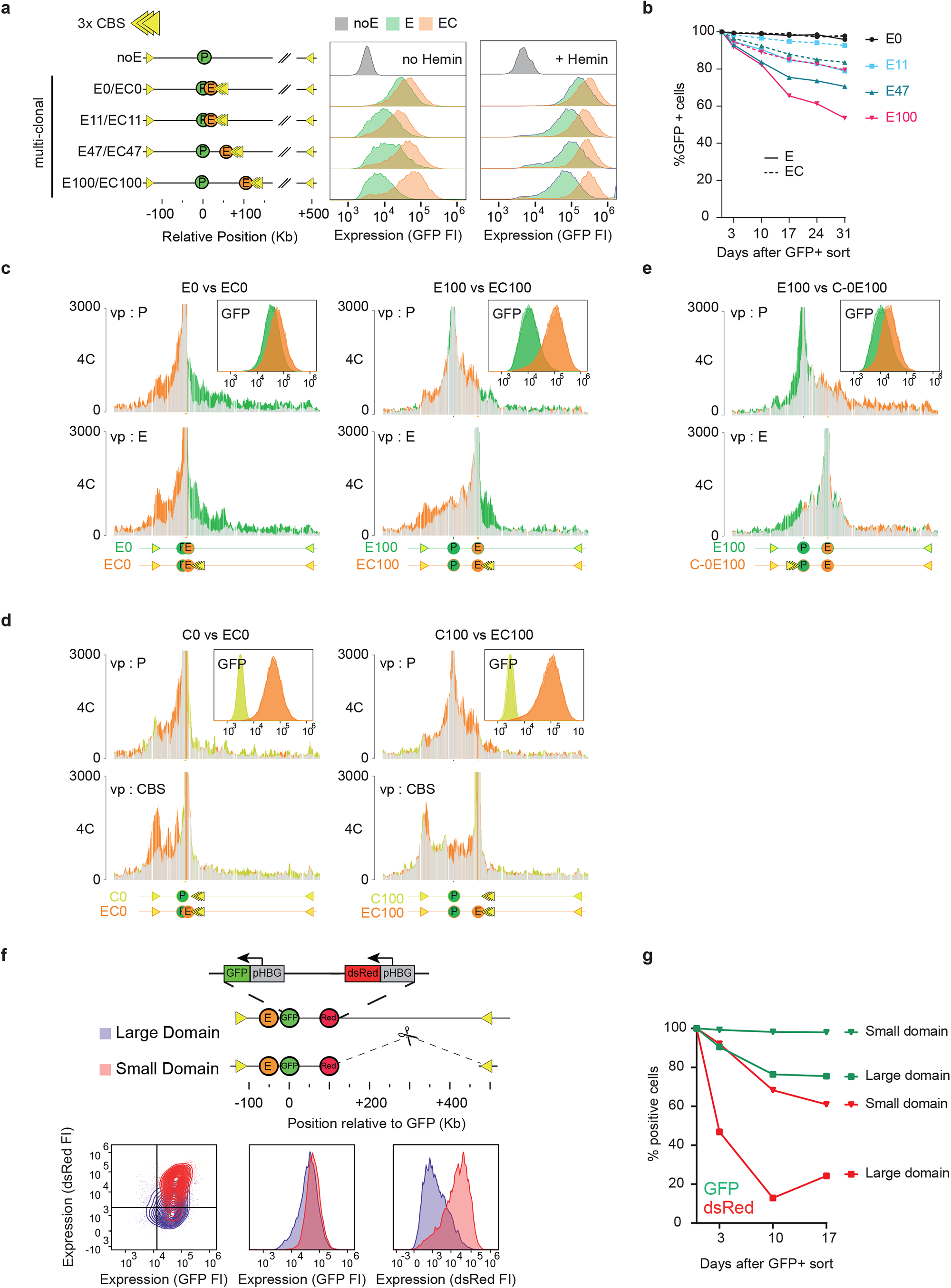
Activated CTCF sites that reduce domain size support enhancer-dependent expression levels and stability. **a**. GFP fluorescent intensity (FI) of multi-clonal (bulk) cell lines carrying the enhancer without (E, in green) and with (EC, in orange) the co-integrated 3x-hCTCF cassette at the indicated positions, cultured without hemin (left panel) or with hemin (right panel). Yellow triangles symbolize the inserted 3x-hCTCF binding sites (CBS). **b**. Percentage of remaining GFP positive cells in multi-clonal (bulk) cell populations carrying the enhancer without (solid line) or with flanking 3xhCTCF (dashed line) at the indicated distances from the reporter gene. At each time point, cells were first treated with hemin for two days prior to FACS analysis. **c**. 4C-seq contact profile overlays, comparing contacts of the integrated gene promoter (P) and µLCR enhancer (E) in the E0 versus EC0 cell line, highlighting in orange their gained contacts and in green their lost contacts due to the presence of the 3x-hCTCF binding sites (CBS). Y-axis: 4C coverage normalized per 1 million *cis*-reads. n= two technical replicates/clone. VP: 4C-seq viewpoint. Also shown at the top right is the overlay of their FACS profiles, demonstrating that at position 0, the 3x-hCTCF cassette mildly supports enhancer-mediated GFP expression. **d**. 4C-seq contact profile overlays comparing contacts of the integrated gene promoter (P, top) and 3x-hCTCF cassette (CBS, bottom) in the EC0 versus C0 cell line (left) and the EC100 versus C100 cell line (right), with contacts gained and lost due to the presence of the enhancer shown in orange and yellow, respectively. Y-axis: 4C coverage normalized per 1 million *cis*-reads. n= two technical replicates/clone. VP: 4C-seq viewpoint. Top right plots show the overlay of their FACS profiles, demonstrating that both in C0 and in C100, the 3x-hCTCF cassette itself does not activate GFP expression. **e**. 4C-seq contact profile overlays comparing contacts of the integrated gene promoter (P, top) and enhancer (E, bottom) in the C-0E100 versus E100 cell line, with contacts gained and lost due to the presence of the downstream 3x-hCTCF sites shown in orange and green, respectively. Y-axis: 4C coverage normalized per 1 million *cis*-reads. n= two technical replicates/clone. VP: 4C-seq viewpoint. Top right plots shows the overlay of their FACS profiles, demonstrating that the downstream 3x-hCTCF cassette only mildly supports enhancer-mediated GFP expression. **f**. Schematic representation of the two dual reporter lines. The µLCR enhancer was integrated 50 Kb downstream of the GFP reporter gene, while a dsRed reporter gene driven by the same HBG1 promoter was integrated 100 Kb upstream of the GFP reporter gene. To place the genes and enhancer in a smaller domain, the 300 kb region between the dsRed reporter gene and the right domain boundary was deleted. Plots show hemin-induced expression (fluorescent intensity, FI) of both genes measured by FACS in the large domain cell line (purple) and the small domain cell line (red), three days after sorting each line for double positive (GFP+/dsRed+) cells. Lower left panel highlights that in the context of a small domain, more cells manage to keep both gene active, and expressed at higher levels. Middle panel shows GFP expression (FI) for both clones and right panel shows dsRed expression (FI) for both clones, highlighting that particularly the expression of the distal dsRed reporter gene benefits from being part of a smaller domain. **g**. The small domain protects against gene silencing, particularly of the distal dsRed reporter gene. Percentage of remaining GFP positive (green lines) and dsRed positive cells (red lines) after sorting the ‘small domain’ cells (triangles) and the ‘large domain’ cells (squares) each for double positive (GFP+/dsRed+) cells and culturing them for the indicated period. At each time point, cells were first treated with hemin for two days prior to FACS analysis.

### Tissue-specific enhancer recruits cohesin to create self-interacting domains and activate CTCF sites for looping

The formation of contact domains relies on cohesin and the degree of self-interaction is believed to reflect the local loop extrusion activity^12,19^. Tissue-specific enhancers have been proposed to recruit cohesin^20,24,26,27,29-32^. To directly test whether indeed our enhancer recruited cohesin to induce intra-domain chromatin interactions, we CRIPSRi depleted cohesin components SMC1A and RAD21 in the E100, EC100 and E407 clones. In all clones, following depletion, we observed a loss specifically of the enhancer-induced chromatin contacts (**Fig. 4a-d and Extended Data Fig. 5**), strongly supporting the hypothesis. To further validate whether the enhancer recruited cohesin to the locus, we performed qPCR-based chromatin immunoprecipitation (ChIP-qPCR) for SMC1A and analyzed cohesin levels at selected sites. In all E-lines with enhancers at varying distances from the reporter gene, except for the E407 line, we found more cohesin deposited at the reporter gene promoter than in the control line lacking an enhancer (**Fig. 4e**). Some increase of cohesin at the promoter was also observed upon proximal (C0) and distal (C100) integration of the 3x-hCTCF cassette without the μLCR (**Fig. 4f**). Cohesin deposition at the promoter was highest in the EC0 and EC100 lines, having both the enhancer and CTCF sites co-integrated at 0 and 100 kb, respectively (**Fig. 4f**). The same observation was made for the integrated CTCF sites themselves: they already recruited cohesin in the C-lines, but cohesin accumulated to higher levels in both EC-lines, having also the enhancer integrated near these CBS (**Fig. 4g**). We then asked whether the enhancer also accumulated cohesin on endogenous CTCF sites. We focused on the EC0 and EC100 cells that showed a strong, enhancer-stimulated chromatin loop between the integrated 3xCTCF cassette and the convergent endogenous CTCF site that we termed the ‘left boundary site’ (**Fig. 3d**). Since this CTCF site is also present on the untargeted copies of chromosome 18, exclusive analysis of its characteristics on the targeted allele was not possible. Despite this technical limitation, 4C-seq directed to this position confirmed that this normally unengaged CTCF site now formed a novel chromatin loop with the integrated CTCF sites in both cell lines (**Extended Data Fig. 4a**). Also, ChIP-qPCR demonstrated that the integrated enhancer deposited cohesin at this left boundary site (**Fig. 4h**). Collectively, from these chromatin topology studies in cohesin-depleted cells and the ChIP-qPCR results, we concluded that the enhancer recruited extruding cohesin complexes to the locus and deposited them at flanking convergent CTCF sites. To further investigate the domain-forming capacity of the enhancer, we created two other cell lines, again through a series of subsequent genetic modifications, with an identically modified CTCF binding landscape, but with a differently located enhancer. For this, the 3x-hCTCF cassette was placed at position 100 (C100), as before, while another CTCF cassette, 3x-mCTCF, with three strong CTCF binding sites selected from the mouse genome^47^ (to have sequences not existing in the human genome), was placed in identical orientation at position 0 (C0). One line then carried the μLCR downstream of C0, while the other cell line had the enhancer downstream of C100. In both scenarios, the left boundary site was activated, judged from increased levels of SMC1A measured by ChIP-qPCR (**Fig. 4i**). When we applied 4C-seq to this site we found that the E0 stimulated its contacts mostly with C0, creating a small domain, while the E100 enabled the left boundary to select the distal C100 as its preferred contacting partner (**Extended Data Fig. 4b**). Also, despite the cell lines having identical CTCF binding landscapes, the distally integrated enhancer (E100) activated the reporter gene to probe a much larger domain (**Fig. 4i**). Thus, the location of the enhancer determined the domain that was formed. The data further suggested that cohesin loaded at the E100 enhancer could readily bypass the ectopic, divergently oriented CBS at C0. In agreement, we observed that cohesin levels at CO were higher with the enhancer at E0 than at E100, while the opposite was true for cohesin levels at C100 (**Fig. 4i**). Our data therefore demonstrated that the tissue-specific enhancer created local self-interacting domains and CTCF-mediated chromatin loops through the recruitment of extruding cohesin complexes. The location of the enhancer dictated which flanking convergent CTCF sites were selected for cohesin stalling and for the formation of domain-spanning chromatin loops.

**Fig 4.**
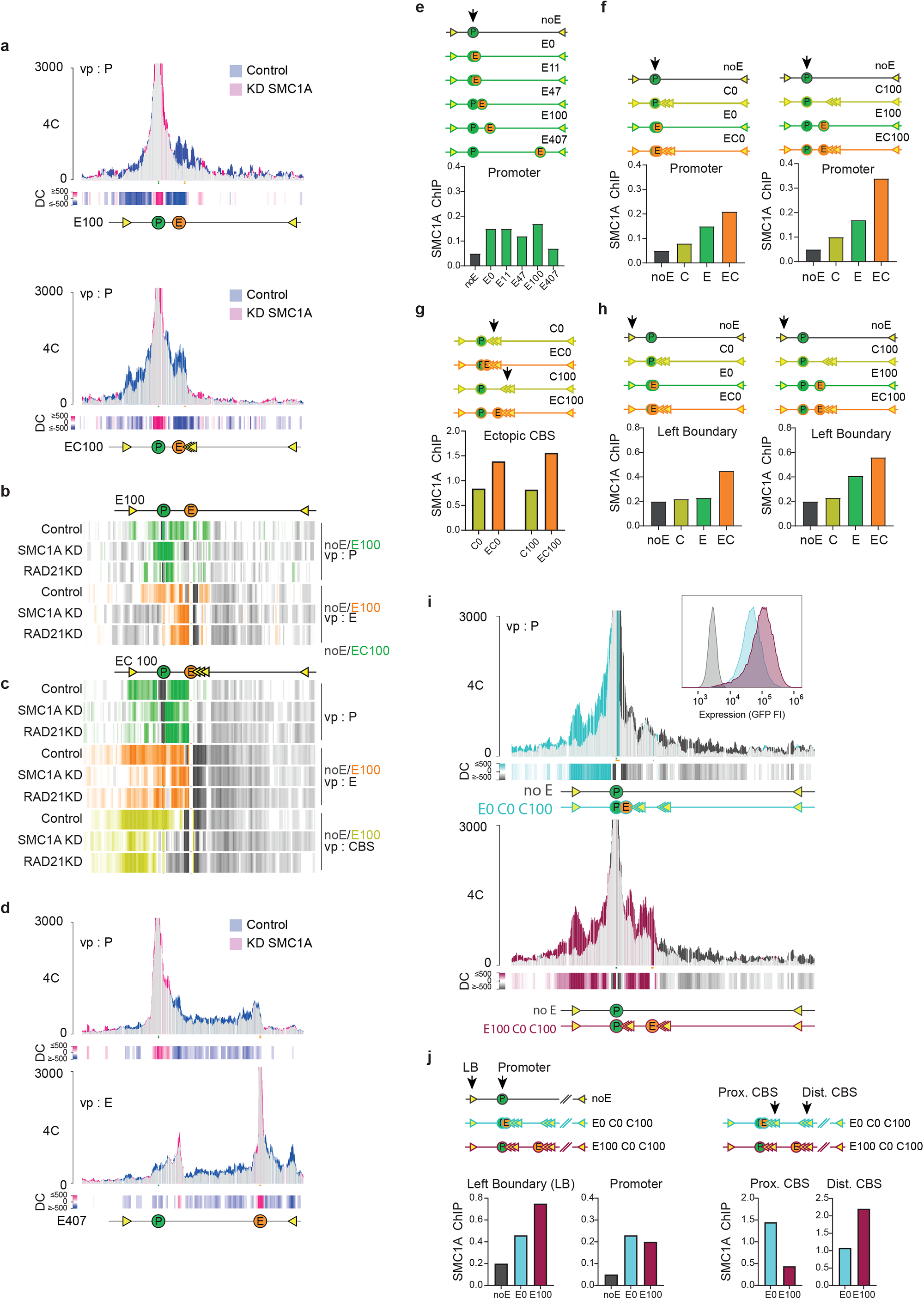
Tissue-specific enhancer recruits cohesin to create self-interacting domains and activate CTCF sites. **a**. SMC1A depletion causes loss of enhancer-induced contacts. 4C-seq contact profile overlays comparing contacts of the integrated promoter (P) in two cell lines, E100 (top) and EC100 (bottom), following CRISPRi depletion of SMC1A (pink) versus contacts measured in cells treated with control sgRNA (blue) (blue). Shared contacts are in gray. Y-axis: 4C coverage normalized per 1 million *cis*-reads. Differential contacts (DC) plotted per 5kb genomic bins are shown below. **b-c**. 4C-seq differential contacts (DC) tracks showing, for cell lines E100 (**b**) and EC100 (**c**), in green the gained contacts of the gene promoter in control cells, SMC1A depleted cells and RAD21 depleted cells, as compared to its contacts in the cell line lacking the enhancer (noE). Similarly, in respectively orange and yellow are shown the gained contacts by the enhancer and the 3x-hCTCF sites in control cells, SMC1A depleted cells and RAD21 depleted cells, as compared to the contacts of their corresponding endogenous chromosomal position in the cell line lacking the enhancer (noE). Note that in all instances, the enhancer-stimulated contacts seen in control cells are (partially) lost again upon cohesin depletion. **d**. SMC1A depletion causes loss of enhancer-induced contacts in E407. 4C-seq contact profile overlays comparing contacts of the integrated promoter (P, top) and enhancer (E, bottom) in E407 cells following CRISPRi depletion of SMC1A (pink), versus those measured in E407 cells treated with control sgRNA (blue) (blue). Y-axis: 4C coverage normalized per 1 million *cis*-reads. Differential contacts (DC) plotted per 5kb genomic bins are shown below. **e-h**. The enhancer deposits cohesin at flanking CTCF sites and the gene promoter. ChIP-qPCR results showing relative enrichment of SMC1A protein bound to the integrated GFP reporter gene promoter (**e, f**), 3x-hCTCF binding sites (**g**) and left boundary (**h**), for indicated E-lines, C-lines and EC-lines. Enrichment values on the y-axis are relative to that measured at a strong SMC1A binding site in K562 cells (see methods). **i**. The location of the enhancer dictates the domain contacted by the gene. Top: 4C-seq contact profile overlay showing the contacts of the integrated reporter gene promoter (P) gained (in blue) when located in the indicated CTCF binding landscape with an immediate proximal enhancer (E0). Bottom: 4C-seq contact profile overlay showing the contacts of the integrated reporter gene promoter (P) gained (in red) when located in the same CTCF binding landscape, but with the enhancer at position E100. In both plots, contacts are compared to that of the promoter in a cell line (noE) lacking integrated enhancers or CTCF sites. Y-axis: 4C coverage normalized per 1 million *cis*-reads. Differential contacts (DC) plotted per 5kb genomic bins are shown below. n= two technical replicates/clone. **j**. The enhancer selects which CTCF sites are activated. ChIP-qPCR enrichment for SMC1A at the left boundary, promoter, proximal integrated 3x-mCTCF binding sites and distal integrated 3x-hCTCF binding sites for noE (dark grey), E0 C0 C100 (blue) and E100 C0 C100 (red) cells.

### Cohesin is required for long-range gene activation but dispensable for short-range enhancer action

Knowing that the enhancer recruits cohesin to mobilize the gene, we then studied cohesin’s requirement for enhancer-mediated gene activation. We took all E-lines with the enhancer at varying distances from the target gene. In the distal configurations (E100 and E407), knockdown of all three cohesin subunits RAD21, SMC3 and SMC1A led to strong down-regulation of the reporter gene expression (**Fig. 5a, b**). At E47, knockdown of the three cohesin factors clearly had a less negative impact on expression, while at the two most proximal E-P combinations, E0 and E11, knockdown of each of these cohesin factors had an entirely opposite, namely positive effect on gene expression. To relate this to the impact of other factors we also knocked-down GATA1, a transcription factor known to support transcription and looping in the β-globin locus^48^, and two components (MED1 and MED21) of mediator^49^, a protein complex important for enhancer function. Different from cohesin, GATA1 and mediator were required for transcriptional activation by all enhancers, irrespective of their distance to the promoter (**Fig. 5a, b**). Thus, the enhancer strictly needed transcription factors and mediator for its activity, but relied on cohesin only for the activation of distal genes, not for the activation of proximal genes. This strongly suggested that an enhancer requires cohesin loop extrusion to bring a distal gene in proximity in order to activate it.

**Fig 5.**
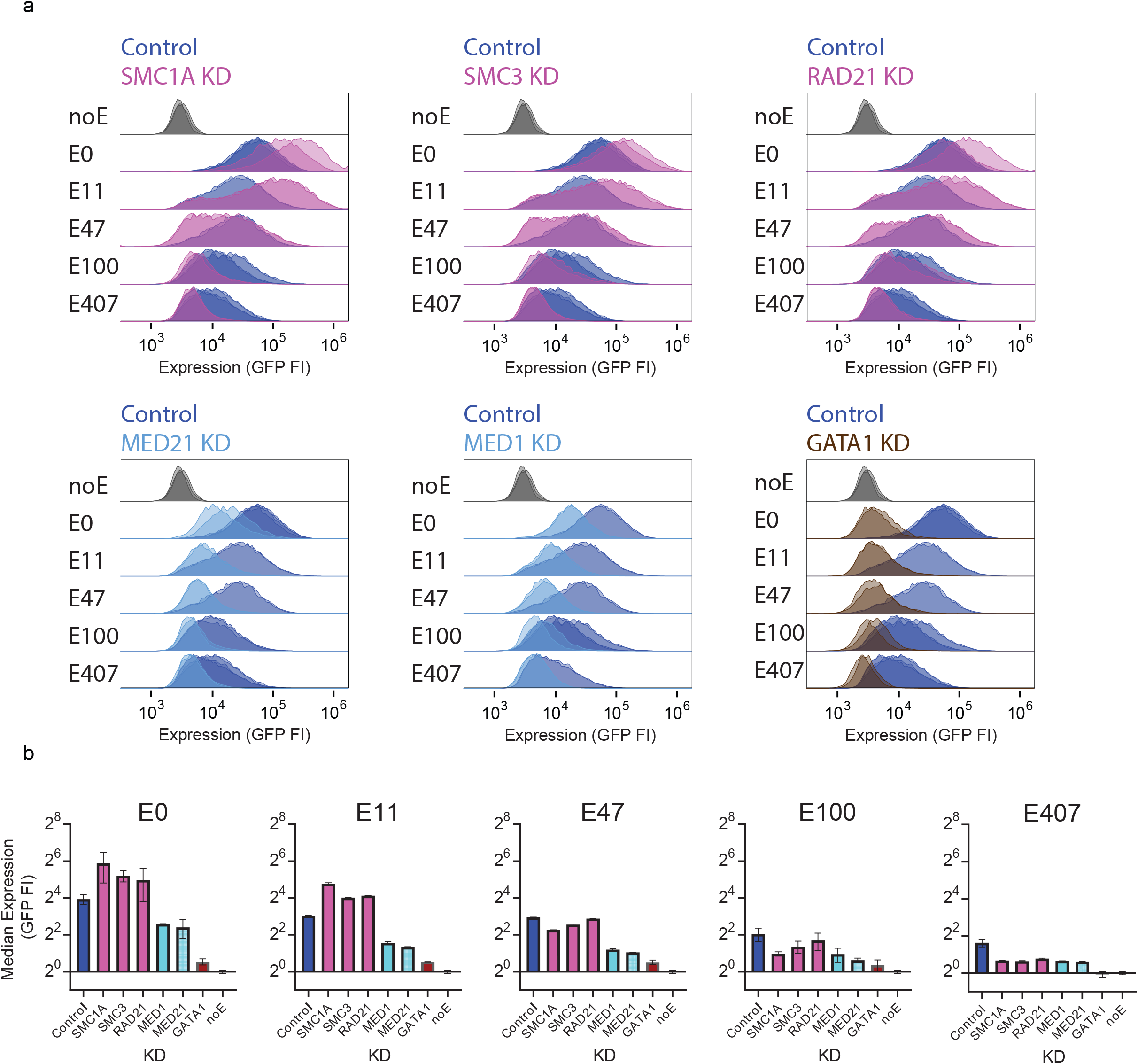
Cohesin is required for long-range gene activation but dispensable for short-range enhancer action. **a**. GFP expression measured by FACS (FI) in arbitrary units (a.u.) in E0, E11, E47, E100 and E407 cells CRISPRi depleted for the cohesin subunits SMC1A, SMC3 and RAD21 (top, in pink), the mediator components MED21 and MED1 (bottom left, in light blue) and the transcription factor GATA1 (bottom right, in brown) as compared to GFP expression in cells treated with a control sgRNA (all panels, dark blue). For comparison, top rows show GFP expression in the cell line lacking an integrated enhancer (noE cells: dark gray). All histograms have at least two replicates. **b**. Median GFP expression measured by FACS (FI) in the different cell lines upon CRIPSRi-mediated knockdown of cohesin components SMC1A, RAD21 and SMC3 (pink), mediator components MED21 and MED1 (light blue) and transcription factor GATA1 (brown) or control (dark blue), plotted relative to the median expression measured in noE cells. Error bars represent standard deviation (SD) of at least two replicates. Note that independent of its distance to the reporter gene promoter, enhancer activity is severely affected upon loss of MED21, MED1 and GATA1, while cohesin is exclusively required for long-range enhancer action, not for activation of an immediately proximal gene promoter (E0).

## DISCUSSION

Enhancers are tissue-specific regulatory DNA elements: they rely on the availability and recruitment of tissue-specific transcription factors for their ability to activate transcription of surrounding genes. By using a unique, bottom-up approach of building many different regulatory landscapes in an inactive chromatin environment, we provide experimental evidence for an emerging model^20,24,26,27,31,50,51^ in which the concerted action of tissue-specific transcription factors at enhancers serves two purposes: (1) they enable recruitment of co-factors, together with which they can stimulate initiation and elongation of the transcriptional machinery at gene promoters and (2) they enable recruitment of cohesin, which, presumably through DNA extrusion, locally stimulates looping and contacts with and between more distal sequences, to form contact domains. We here show that cohesin recruitment is necessary for enhancers to activate distant, but not proximal, genes. Since cohesin forms chromatin loops, this supports the debated idea that long-range gene activation, certainly in an inactive chromatin context, requires chromatin looping. Importantly, developmental genes which rely on distant tissue-specific enhancers for their activation, often locate near gene deserts or in gene-poor segments, being parts of the genome that by default are inactive. Interestingly, our E-P pair functioned perfectly well (even better) in cells with reduced cohesin not only when they were immediately juxtaposed in E0, but also when they were separated by 11kb (E11). Even when 47kb apart (E47), the pair was relatively resistant to reduced cohesin levels. This, and similar observations made in a parallel study on the *Shh* locus^52^, argues that we may need to better define the meaning of ‘long-range’ when discussing mechanisms of gene activation. We postulate that the type (strength) of the enhancer and the genomic context determines whether the enhancer relies on cohesin-dependent or –independent mechanisms^25,28^ to fully activate a target gene over intermediate distances (let’s say 0-100kb). Molecular condensates formed through non-specific interactions between intrinsically disordered domains of enhancer- and promoter-associated transcription factors and Mediator may enable cohesin-independent E-P communication over such distances^53-56^. Over further distances, we expect that enhancers increasingly also need cohesin for gene activation.

We further find that the genomic location of the enhancer determines which flanking pair of convergent CTCF sites is selected as boundaries of the contact domain. If they encompass a small domain, they can support the enhancer in conferring strong expression to the target gene and protect it from silencing. We propose this is a consequence of concentrated loop extrusion activity. We, like others^36,45,46^, find that transcriptional activity is controlled by more than just E-P contact frequencies. Our data further showed that the enhancer stimulated the distal gene not only in E-P contacts, but also in contacts with intervening and immediately surrounding sequencing. This may suggest that recruited cohesin does not stay anchored at the enhancer, but extrudes and simultaneously migrates away from its entry site. Interestingly, modeling showed that local chromatin loops can stimulate enhancer-promoter communication, even if they are not directly anchored at the regulatory elements^57^. In summary, our work demonstrates that an enhancer can recruit cohesin to stimulate the formation of a self-interacting domain, engage flanking CTCF sites in loop engagement and activate expression of distal target genes.

## METHODS

### Cell culture

Human erythroleukemia K562 cells were grown at 37°C at 5% CO2 in RPMI 1640 (Gibco) with 10% FBS (Sigma) and 1% Penicillin-Streptomycin (Gibco). Density was kept between 1×10^5^ and 5×10^5^ cells per ml medium. Cells were routinely tested for mycoplasma. Prior to every experiment, cells were weekly bulk sorted for two consecutive weeks on Becton Dickinson SORP FACSAria FUSION Flow Cytometer. Gating was based on noE cells or Fluorescence minus one controls. Flow cytometry analysis was done three days after sorting on a Beckman Coulter Cytoflex S. For differentiation, culture medium was supplemented with porcine hemin (Sigma, 30 µM final concentration) one day after sorting. Hemin and medium were refreshed one day later. 4 mM hemin stock solutions were prepared according to^58^ and kept at -20°C. For long term culturing and FI monitoring, at each indicated time point an aliquot of cells was taken, treated with hemin for two days and FI monitored. To obtain irreversibly silenced populations of cells, GFP negative cells where weekly sorted for six (E100) or nine (E407) consecutive weeks.

### Targeting constructs

Targeting constructs contained one or two regulatory elements (µLCR^41^, µLCR::3xmCTCF^47^, µLCR::3xhCTCF (described below), dsRed::pHBG^59^) of interest flanked by two ∼1Kb-sized site-specific homology arms that were amplified from genomic DNA with primers as indicated in **Supplementary Table 1**. Fragments were combined into a plasmid by In-Fusion Cloning (Takara).

The three human CTCF sites (3xhCTCF) where selected from^60^ and PCR-amplified from the following genomic DNA positions: chr19:41650330-41650595, chr7:39559582-39559824 and chr13:21498993-21499294, and combined by overlap-extension PCR using primers as indicated in **Supplementary Table 1**.

### Clonal Cell line generation

The founder cell line containing the d2eGFP reporter driven by human pHBG1 and µLCR enhancer was generated by Tol2 transposition. The founder µLCR is flanked by sleeping beauty (SB) terminal inverted repeats (ITRs) and LoxP sites which split a puromycin gene driven by pSV40 such that it is not functional. GFP-expressing cells upon transgene transposition were single-cell sorted and its integration sites were mapped using TLA^61^. The µLCR enhancer was removed by transient transfection of a CRE-recombinase encoding plasmid. The clone that had the transgene integrated at chromosome 18 (position: 19609009) was selected for further experiments and designated “noE”.

Regulatory elements where integrated in target cell lines using CRISPR/Cas9 and homology directed repair. Cas9 plasmid (pSpCas9(BB)-2A-BFP (a modified version of PX458 (Addgene plasmid # 48138), in which eGFP is replaced for tagBFP)) containing a position-specific single-guide was co-transfected with a targeting plasmid and GFP expressing cells were single-cell sorted using flow cytometry, expanded, genotyped and sanger-sequenced for correct integration of the transgene. Guide sequences specific for position of interest were cloned between BbsI sites and are listed in **Supplementary Table 2**.

The µLCR enhancer was targeted to noE cell line to positions 11 Kb and 47 kb upstream of eGFP-reporter to obtain the respective clonal cell lines. µLCR inserted at position 407 kb carried an additional Anch4 imaging platform (kind gift from K. Bystricky). µLCR::3xhCTCF was targeted to positions 0 Kb (pos0) and 100 Kb (pos100) to obtain EC0 and EC100, with loxP sites flanking the 3x-hCTCF cassette and FRT sites flanking the μLCR. Additionally, µLCR::3xmCTCF was targeted to noE to obtain EmC0. E0 and E100 were generated by transient transfection of a CRE-recombinase (Cre) encoding plasmid. C0, mC0 and C100 were made by expressing the recombinase Flippase (Flp) in the respective EC-lines. E100 C0 C100 clonal cell line was generated by targeting µLCR::3xhCTCF to pos100 in cell line mC0. For E0 C0 C100, µLCR::3xmCTCF was targeted to C100, pos0. C-0 E100 was generated by deleting the 3’ part of eGFP and subsequently repairing it with a targeting construct containing 3xhCTCF::eGFP flanked by homology arms. Homology arm primers and genotyping primers are indicated in **Supplementary Table 1**.

To place the E407 enhancer immediately upstream of the GFP reporter gene, the genomic region between the integrated eGFP and the integrated μLCR at position 407 kb was removed by transient transfection of two Cas9 plasmid (pSpCas9(BB)-2A-BFP plasmids each containing a specific single guide targeting the upstream or downstream sites of this region. Guides are listed in **Supplementary Table 2**.

Double reporter cell lines where generated by targeting dsRed::pHBG1 to pos100 in a separately generated E50 cell line that also had a flanking Anch4 sequence. GFP/dsRed double positive cells were selected to obtain single clones. The intervening region between dsRed reporter and right boundary was removed as described above.

### Multi-clonal cell line generation

µLCR or µLCR::3xhCTCF were targeted as for the clonal cell lines on the same day to noE cell line at positions 0 Kb, 11 Kb, 47 Kb and 100 kb upstream of eGFP-reporter to obtain the respective E- and EC-lines. After five days, GFP-expressing cells were bulk sorted weekly for two consecutive weeks and analyzed in the presence or absence of hemin three days after the second sort. Cells that were not treated with hemin were kept in culture and induced with hemin prior to analysis at days 10, 17, 24 and 31 after the second sort.

### KRAB silencing

For gene knock-down experiments, dead-Cas9 (dCas9) fused to Krüppel-associated box (KRAB) was randomly integrated in the genome of the target cell line via lentiviral transduction of a modified version of pHR-SFFV-dCas9-BFP-KRAB (Addgene plasmid # 46911) carrying a P2A blasticidin selection and UCOE element (kind gift of Marvin Tanenbaum). Top 50% BFP-expressing cells were bulk sorted and kept under 0.7 μg/ml blasticidin selection (Sigma). Single guide RNA (sgRNA) sequences against genes of interest were cloned into lentiviral targeting plasmid pU6-sgRNA EF1Alpha-puro-T2A-BFP (Addgene plasmid # 60955). Optimal target sequences were selected from^62^. dCas9::KRAB containing target cells were first sorted for GFP for two consecutive weeks and transduced with sgRNA-coding virus two days after the second sort. Cells recovered for one day before puromycin selection (Sigma, 1 µg/ml final concentration) was started. Crosslinking for 4C and analysis by flow cytometry was performed 5 days post puromycin selection. sgRNA sequences are listed in **Supplementary Table 2**.

### Flow Cytometry analysis

Flow Cytometry Standard (FCS) files were analysed with FlowJo Software. Fluorescence compensation was applied based on single-fluorophore samples. Cells were gated for live single cells based on FSC-A, SSC-A and FSC-W. For silencing experiments, cells were considered GFP positive based on gating on noE cell line treated with hemin. For KD experiments, cells were considered sgRNA-positive based on control cells containing dCas9::KRAB, but no sgRNA as indicated in **Supplementary Information**.

To calculate relative fluorescence, median GFP FI of KD samples were divided by control KD samples after subtraction of GFP FI for noE sample.

### 4C-seq

4C-seq was performed as described^44^. Eight to ten million cells were cross-linked with formaldehyde. DNA was digested *in situ* with Csp6I (first cutter) and NlaIII (second cutter). Indexed Illumina sequencing adapters were introduced to ligation fragments of interest with a two-step PCR strategy. All viewpoint specific primers are listed in **Supplementary Table 1**. Technical replicates were processed on the same day.

### 4C analysis

FastQ files were demultiplexed and processed as in^44^ (https://github.com/deLaatLab/pipe4C) Reads were mapped against versions of hg19 that were modified to contain the aforementioned insert sequences (modified genomes) at the experimentally validated coordinates.

Plotting and contacts counting was done in R (https://www.R-project.org/). Blind fragments were omitted for analysis. Read counts were then normalized to a million mapped intra chromosomal reads (normalized reads) excluding the two highest covered fragments and 21 fragment end rolling mean scores were calculated for every fragment end. For plotting of overlay profiles of distinct modified genomes, positions of fragment ends were shifted based on the coordinates of the inserted sequences, such that common fragment ends were aligned.

For E-P contact counting, 4C profiles with viewpoint eGFP reporter were used. Reads normalized to 1 million cis-reads were counted for the 11 non-blind enhancer fragment ends that could be uniquely mapped using all modified genomes and averaged. Mean coverage per fragment was used for plotting. For differential contact tracks, normalized reads per fragment were averaged per 5 kb bins for each profile and subtracted, which resulted in a number for the average differential contacts per fragment in the 5 Kb bin. Differential contacts were then represented in a color range, where less than 50 normalized read difference were indicated in white and then color-scaled between 50 and 500 differential contacts, as indicated.

### ChIP-qPCR

For each batch of ChIP experiments 2.5 million cells were sorted for GFP expression. Cells were cultured for four days to obtain at least 25 million cells, followed by fixation for 10 min at 4°C in 1% PFA. From this point onward, cells were processed via the ChIP-IT High Sensitivity kit (Active motif) as per manufacturer’s instructions. Chromatin was sheared to 200-500 bp fragments on a Bioruptor Plus (Diagenode; 5 × 5 cycles of 30 s on and 30 s off, at the highest power setting). Immuno-precipitation was carried out by adding 5 mg of the appropriate antiserum (SMC1: A300-055A, Bethyl) to approx. 30 mg of chromatin and incubating on a rotator overnight at 4°C in the presence of protease inhibitors. Following addition of protein A/G agarose beads and washing, DNA was eluted using DNA purification elution buffer (Active motif). The eluted DNA was used in qPCR using primers targeting putative cohesin-bound positions in the integration site (coordinates) and normalized over a tested cohesin bound site (chr11:4658282-4658362). All primers are listed in **Supplementary Table 1**.

### Public data

Public ChIP-seq and RNA-seq (mapped to human genome release 19) tracks were downloaded from ENCODE portal^63^. The following data-sets from ENCODE were used: K562 SMC3 ChIP-seq (Encode, Michael Snyder, ENCSR000EGW, ENCFF479BWQ), ChIP-seq CTCF ChIP-seq (Encode, Bradley Bernstein, ENCSR000AKO, ENCFF000BWF), K562 RNA-seq (Encode, Brenton Graveley, ENCSR000AEN, ENCFF657EOD, ENCFF578WIM). H3K27ac (Encode, Bradley Bernstein, GEO: GSM733656) and H3K27me3 (Encode, Michael Snyder, GEO: GSM788088).

## Data availability

Raw sequencing data and mapped wig files are available from the Gene Expression Omnibus (GEO) under accession GSE180566.

## Code availability

Code for processing 4C data used in this study can be found at https://github.com/deLaatLab/pipe4C.

## Acknowledgements

We thank Amin Allahyar for help with 4C analysis and de Laat lab members for discussions and feedback. This work is part of the Oncode Institute, and was funded by a VICI grant (724.012.003) from the Netherlands Organisation for Scientific Research (NWO), an NWO Groot grant (2019.012), a Fondation Leducq (14CVD01) Transatlantic Network grant and an EU MSCA-ITN grant (ENHPATHY 860002).

## Author contributions

N.J.R., K.S., P.H.L.K. and W.d.L. conceived experiments; N.J.R., M.J.A.M.V., Y.O., C.v.-Q, A.F., T.F., Z.d.A.d.R, P.H.L.K. generated constructs and cell lines; N.J.R. and S.v.d.E. performed Flow Cytometry; N.J.R. and S.J.D.T. performed KD experiments; N.J.R., S.J.D.T, M.J.A.M.V and T.F. performed 4C experiments;

N.J.R., R.H. and K.S. designed and performed ChIP-seq experiments; N.J.R. and P.H.L.K. performed bioinformatics analyses; N.J.R., K.S., P.H.L.K. and W.d.L. wrote the manuscript with input from other authors.

---

**Extended Data Fig. 1.**
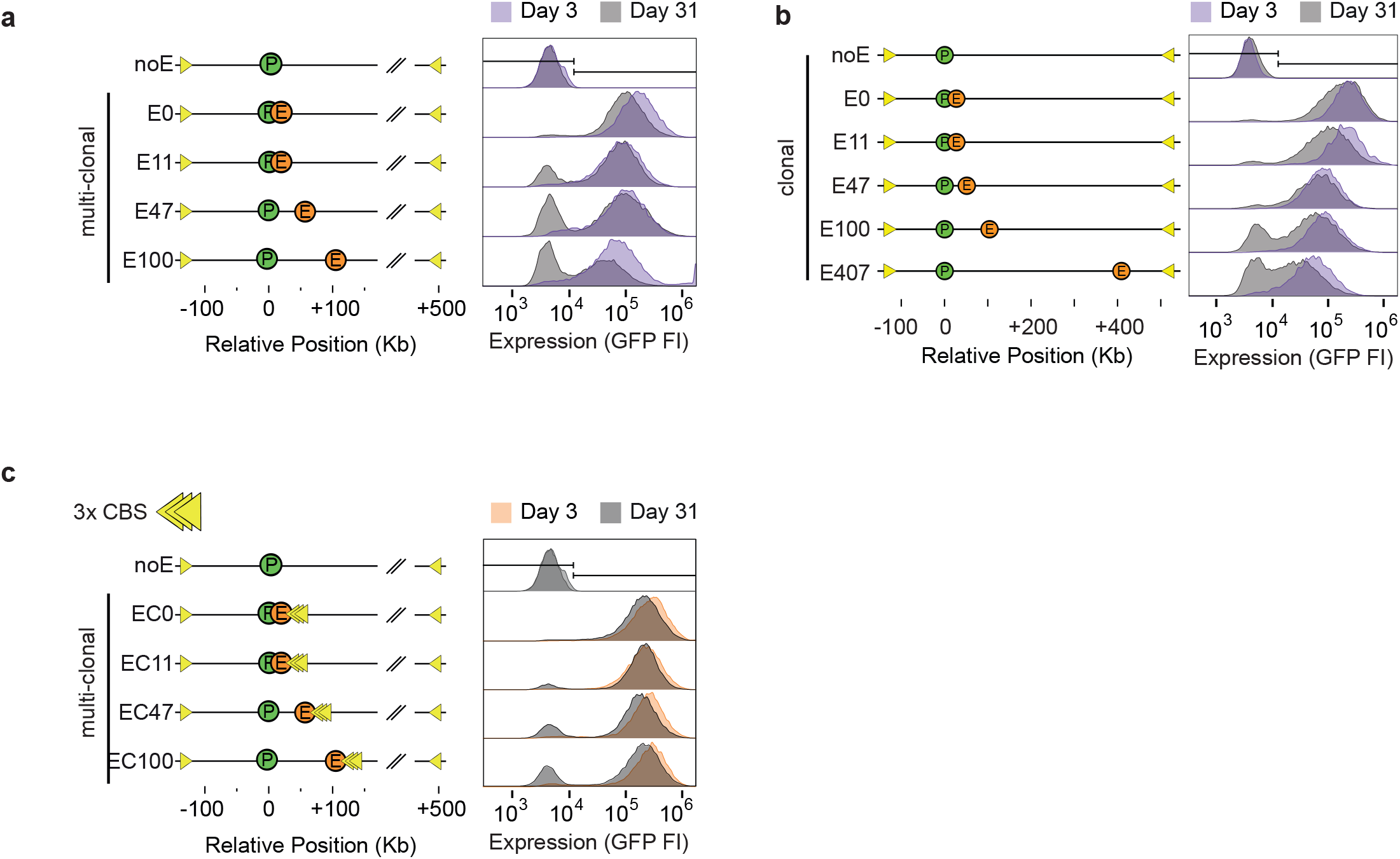
Distance-dependent enhancer protection against gene silencing in long-term cell cultures. **a**. Overlay FACS profiles of multi-clonal cell populations without (noE) or with the enhancer integrated at a given distance (E0-E11-E47-E100kb) from the reporter gene, cultured for respectively 3 and 31 days after sorting for GFP-positive cells. The profiles show that over time, the gene silences with a rate that is related to enhancer distance. **b**. As in **a**, but for clonally selected lines without (noE) or with the enhancer integrated at a given distance (E0-E11-E47-E100kb) from the reporter gene. **c**. As in **a**, but for multi-clonal cell populations without (noE) or with a co-integrated enhancer-CBS at a given distance (EC0-11-47-100).

**Extended Data Fig. 2.**
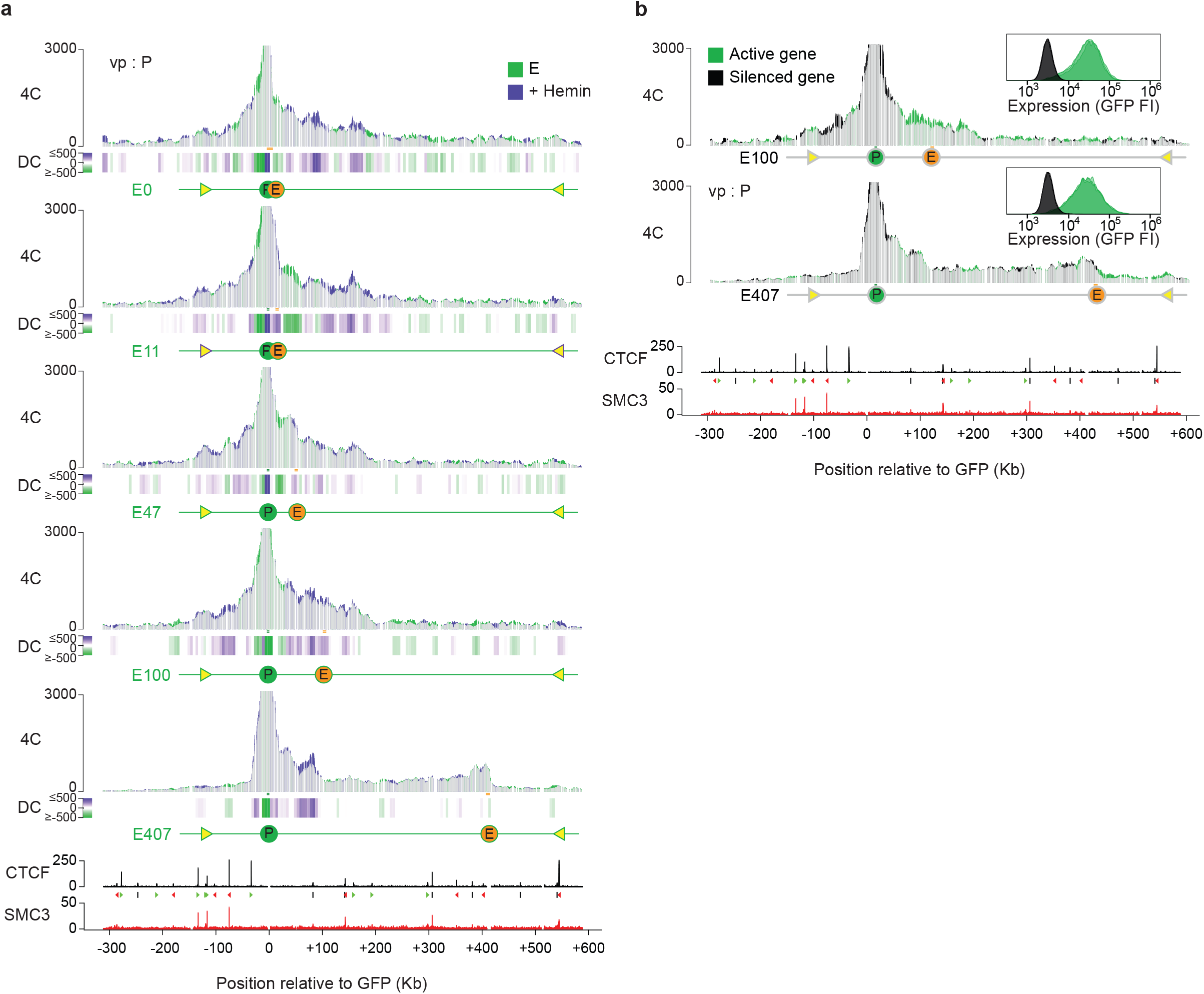
Hemin induced increased expression or reduced expression by prolonged silencing only mildly affects P/E contact frequency. **a**. 4C-seq contact profile overlays comparing contacts of the integrated promoter (P) in the indicated E-lines cultured in the absence (green) or presence of Hemin (purple). Y-axis: 4C coverage normalized to 1 million *cis*-reads. Shared contacts are in light gray. Track below shows the differential contacts (DC) per 5 kb binned fragments. n= two technical replicates/clone. CTCF and SMC3 ChIP-seq signal tracks are shown for reference. Positions are in kb with respect to the integrated GFP reporter gene. **b**. 4C-seq contact profile overlays comparing the contacts of the integrated GFP reporter gene promoter in the expressing (green) and long-term silenced (non-expressing: black) E100 and E407 cell lines. DC-tracks plotted below each overlay. CTCF and SMC3 ChIP-seq signal tracks are shown for reference. Positions are in kb with respect to the integrated GFP reporter gene. GFP expression (FI) for the expressing (GFP+) or long-term silenced cell populations are shown in the top right panel.

**Extended Data Fig. 3.**
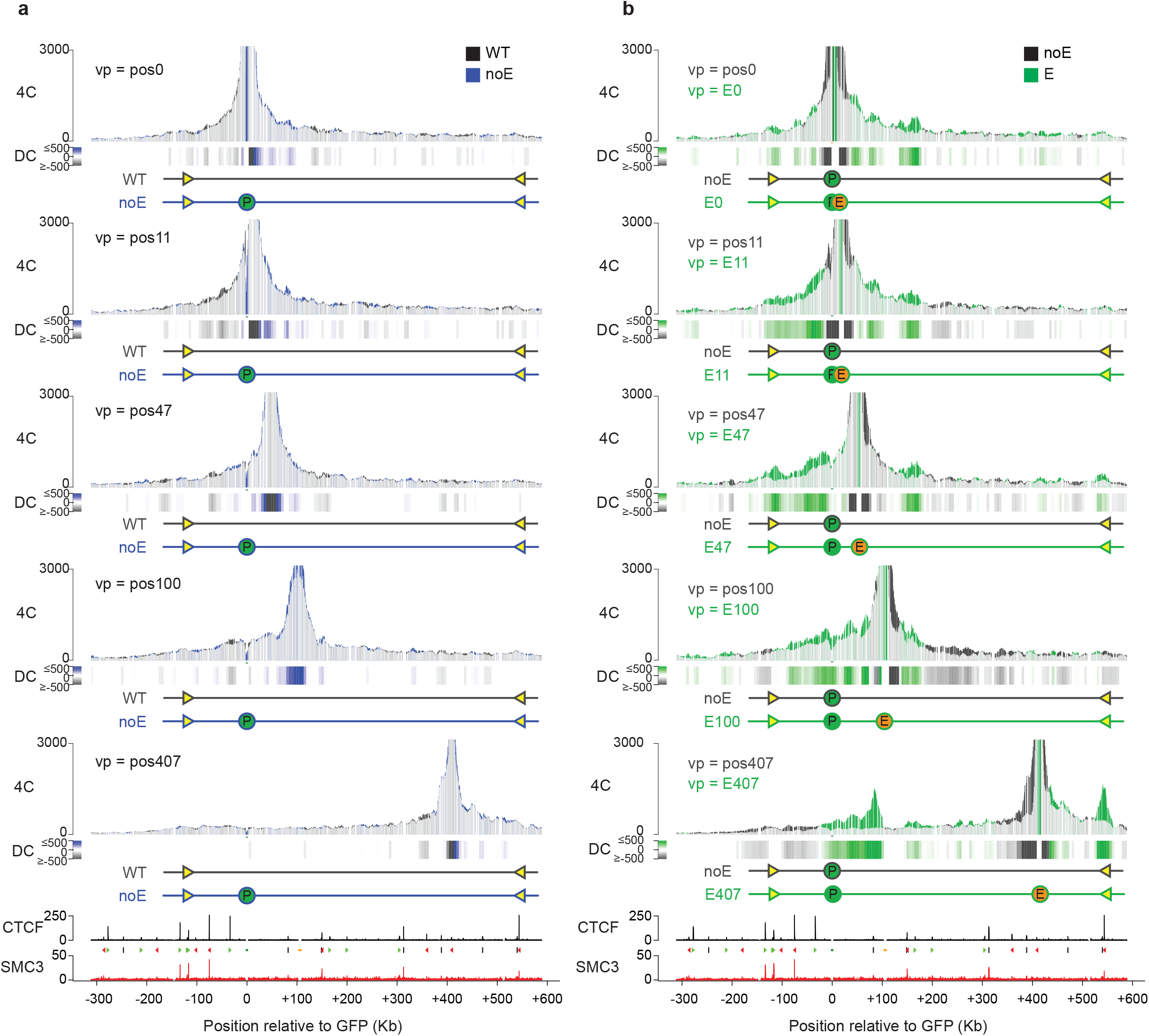
Tissue-specific enhancer, but not the integrated gene cassette, stimulates the formation of local self-interacting chromatin domains. **a**. The integrated gene cassette has little impact on contacts formed by surrounding sequences. 4C-seq contact profile overlays comparing contacts for the endogenous positions (0, 11, 47, 100 and 407 kb upstream of GFP reporter) in noE (blue) and WT (dark grey) cells. Y-axis: 4C coverage normalized per 1 million *cis*-reads. Shared contacts are in light gray. Track below shows the differential contacts (DC) for fragments binned per 5 Kb. n= two technical replicates/clone. **b**. The integrated enhancer engages in new contacts with surrounding sequences. 4C-seq contact profile overlays comparing contacts of the enhancer integrated at the indicated positions, versus contacts of the corresponding endogenous location in cells lacking an integrated enhancer (noE cells). DC-tracks plotted below each overlay. n= two technical replicates/clone.

**Extended Data Fig. 4.**
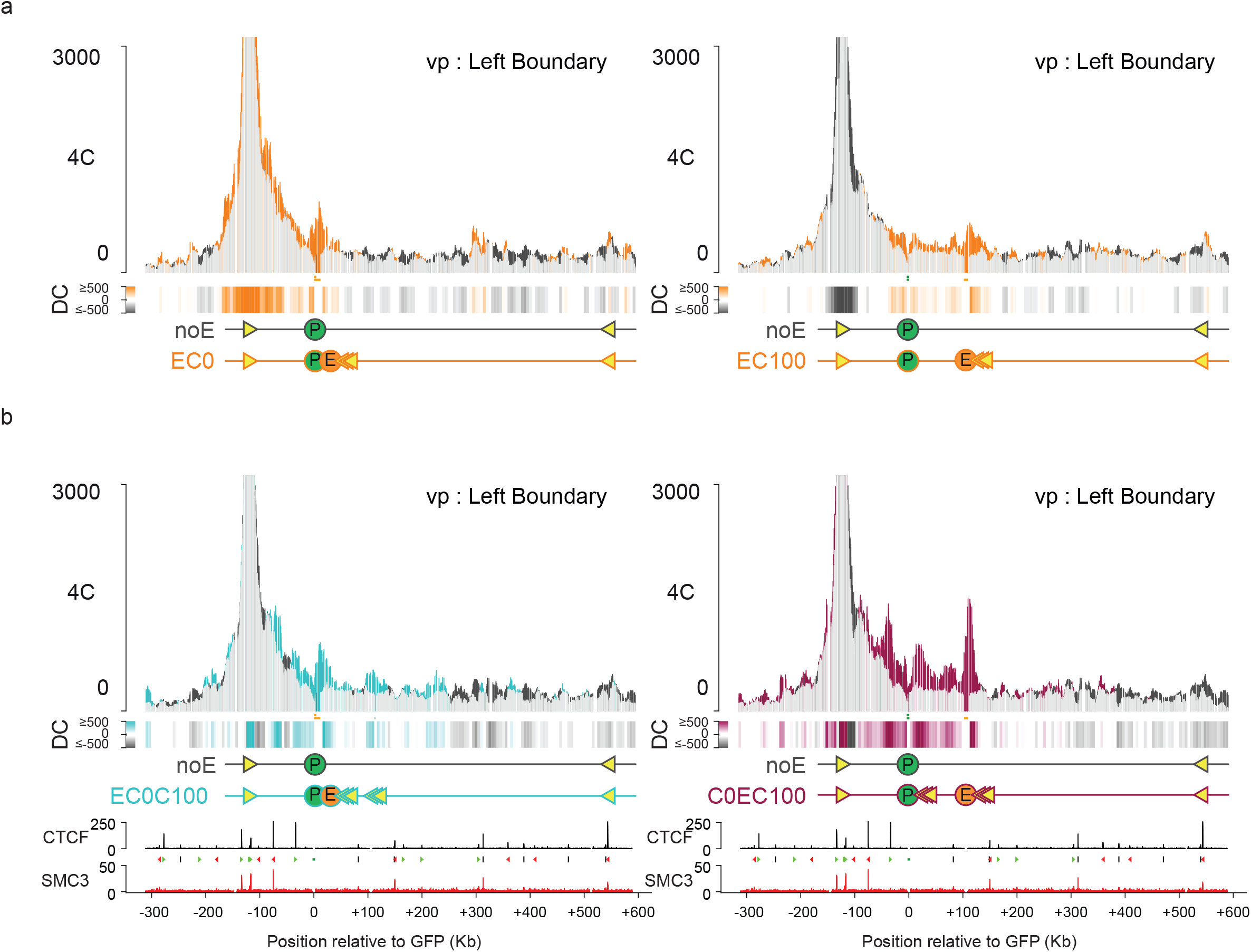
Enhancer selects flanking CTCF sites for engagement in chromatin looping. **a**. 4C-seq contact profile overlays showing contacts gained by the left endogenous boundary in EC lines, in orange, as compared to the left boundary contacts in the cell line lacking an integrated enhancer or integrated CTCF sites (noE: dark gray). Shared contacts are in light gray. y axis: 4C coverage normalized per 1 million *cis*=reads. n= two technical replicates/clone. **b**. 4C-seq profile overlays showing in blue (left) contacts gained by the left endogenous boundary in the cell line having the enhancer at E0 and two CTCF cassettes at C0 and C100, and in red (right), contacts gained by the left endogenous boundary in the cell line having the enhancer at E100 and two CTCF cassettes at C0 and C100. DC-tracks plotted below each overlay. n= two technical replicates/clone.

**Extended Data Fig. 5.**
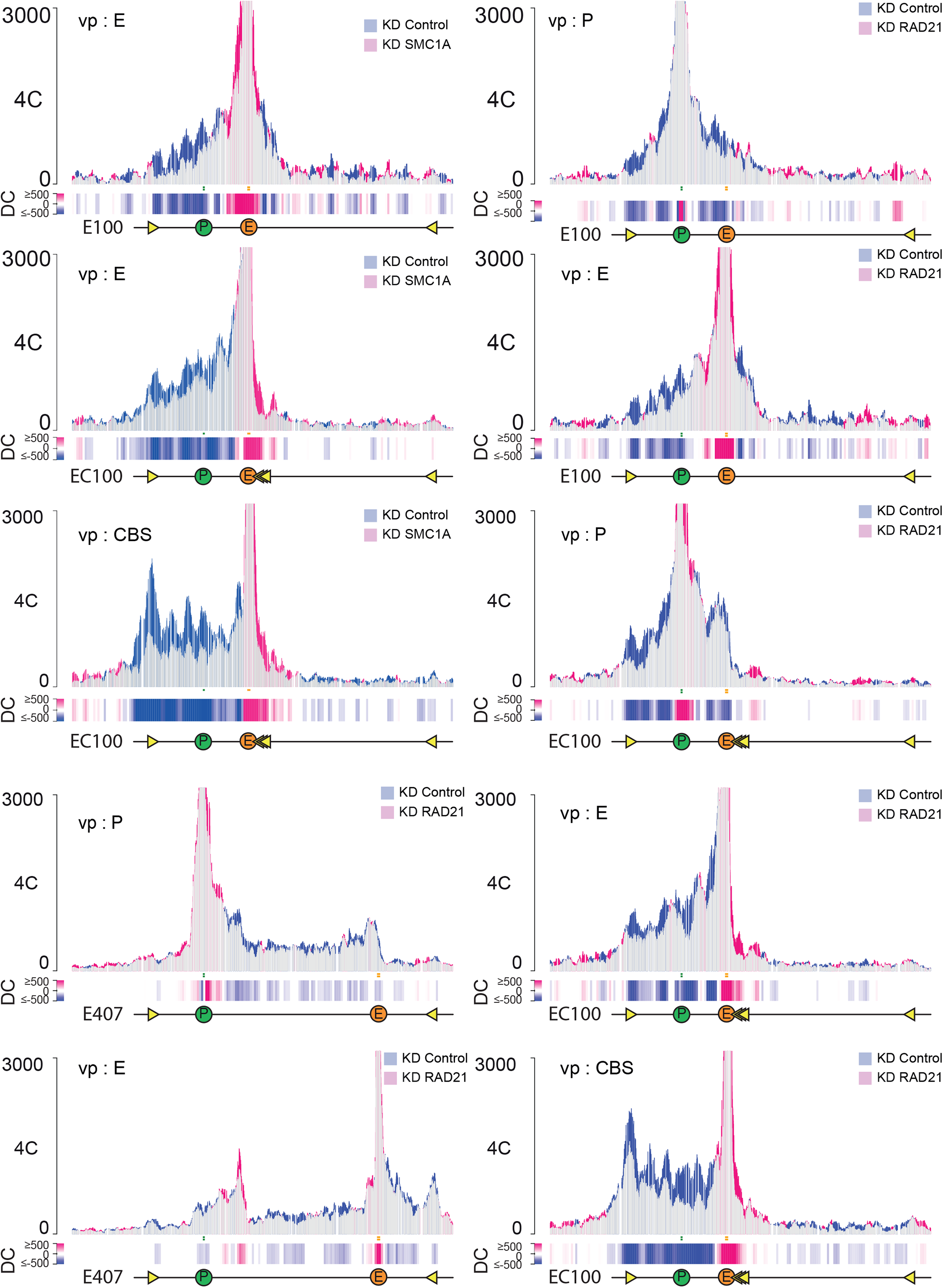
Tissue-specific enhancer relies on cohesin to create self-interacting domains. 4C-seq contact profile overlays comparing contacts of the integrated GFP reporter gene promoter (P), µLCR enhancer (E) and 3x-hCTCF in the E100, EC100 and E407 lines in control (blue) cells and in cells (partially) depleted for RAD21 or SMC1A (pink). Shared contacts are in light gray. Y-axis: 4C coverage normalized to 1 million *cis*-reads.

