## Supplementary Tables for "Building regulatory landscapes: enhancer recruits cohesin to create contact domains, engage CTCF sites and activate distant genes"

**Table S1. PCR Primers Sequences**

*Homology arms targeting constructs*

| | |
|---|---|
| pos-50_Left_HA_FWD | AACACGCTAGGTGTACTGCC |
| pos-50_Left_HA_REV | ACGCACAGCTTTGACAAAAGG |
| pos-50_Right_HA_FWD | GGGGCCCCAGCCCTAGT |
| pos-50_Right_HA_REV | TGAAAGTACCATGGGCGTTGT |
| pos0_Left_HA_FWD | GTGTGTGTGCTGGCTGATGAC |
| pos0_Left_HA_REV | TGCCCGAAGGACCGCGC |
| pos0_Right_HA_FWD | ACGTCGGCGGTGACGGTGAAG |
| pos0_Right_HA_REV | GTCACCGAGCTGCAAGAACTC |
| posMin0_Left_HA_FWD | GCATGGATGTCCAGTGATACA |
| posMin0_Left_HA_REV | TACACCTCCAATTGACTCAAATG |
| posMin0_Right_HA_FWD | AGGTGTACCTGTGGATGTATTT |
| posMin0_Right_HA_REV | AGACCACAGTTTCAGCGCA |
| pos11_Left_HA_FWD | CATGAGAGCCCACCCCTATTC |
| pos11_Left_HA_REV | AGGAGGGGACAGAAAGCAAATC |
| pos11_Right_HA_FWD | CCCCTTTGGTTCAGAATGTCA |
| pos11_Right_HA_REV | TGTACGGACTTCATGCCAGG |
| pos47_Left_HA_FWD | TAGGTCACTCCCTCCATCCTC |
| pos47_Left_HA_REV | CAGGAGCTGACTGGGGACT |
| pos47_Right_HA_FWD | AAGTAGCTACCAACCCAGGC |
| pos47_Right_HA_REV | AACAAAACTTATGAGGTCCATCAAT |
| pos100_Left_HA_FWD | CCAACATTCCCAGAACCAAAC |
| pos100_Left_HA_REV | CTGAAAACGGTGAGGTTTCTG |
| pos100_Right_HA_FWD | GGAGGTTAAATTTTGCCCACT |
| pos100_Right_HA_REV | GGTCCCTAGAGTTCTGTGTCAC |
| pos407_Left_HA_FWD | TTTTCCACCTGCATCTGCCT |
| pos407_Left_HA_REV | ACCCTGTTGTCTACCTAGAGAGG |
| pos407_Right_HA_FWD | TGGGGAAACACTGAGTAATGGT |
| pos407_Right_HA_REV | CCATCCCATGGGCTTGCTAT |
| dsRed_Left_HA_FWD | AGCACTCACATCTTGCCACT |
| dsRed_Left_HA_REV | AGGGGCCCACTGAAGAG |
| dsRed_Right_HA_FWD | AGGGGAGGAAGTGAGAGACA |
| dsRed_Right_HA_REV | GTTCAGCATGGACTAGGGGG |

*human 3x CTCF*

| | |
|---|---|
| CTCF_STAM_A2_FWD | AGAGCGAGATTCCGTCTCAA |
| CTCF_STAM_A2_REV | AGGACAAGCAACAATGGCTGGCCCATAGTA |
| CTCF_STAM_A4_FWD | TGGGCCAGCCATTGTTGCTTGTCCTTCCTTCCTGT |
| CTCF_STAM_A4_REV | CCTGCAAACTGAACTCCTGACCCCTCACAA |
| CTCF_STAM_E2_FWD | TCAGGAGTTCAGTTTGCAGGTGGCTTGACT |
| CTCF_STAM_E2_REV | TTTGATTTCCTTCACTCTGGAA |

*Genotyping primers*

| | |
|---|---|
| uLCR_FWD | CAAGGTTTCACTCTGCTGTC |
| uLCR_REV | TGAATGAGGCTTGAGTACAG |
| 3xhCTCF_FWD | TCTAGATTAGACATAGGCAAGCACA |
| 3xhCTCF_REV | TCTGCCACTGCCTAGTTGAG |
| 3xmCTCF_FWD | TGTCCAACAGAAGATCTTACCACA |
| 3xmCTCF_REV | CTAGTGTGGGCATCGTCCAG |

| | |
|---|---|
| dsRed_FWD | CCCCTAAGCTATCAGGTTGATTGA |
| dsRed_REV | AGCAATAGCATCACAAATTTCACA |
| pos-50_Left_HA_FWD | GCATTATTCATAATAGCACATCTCAATTC |
| pos-50_Right_HA_REV | TGGCACAGCTATATCTAAGGCG |
| pos-50_noInsert_FWD | GCATTATTCATAATAGCACATCTCAATTC |
| pos-50_noInsert_REV | TGGCACAGCTATATCTAAGGCG |
| posMin0_Left_HA_FWD | CAGGACAGGGGATGAGCTT |
| posMin0_noInsert_FWD | TCCGAGAATTCCAGAAAATGATG |
| posMin0_noInsert_REV | ACCTCCCCCTGAACCTGAA |
| pos0_Left_HA_FWD | GTATTTACCACTGGATAAGTG |
| pos0_Right_HA_REV | AGCTTACCATGACCGAGTAC |
| Pos0_noInsert_FWD | TCCTGTGTAAAGCTGGATCC |
| Pos0_noInsert_REV | ACTTCAACTGTAGGCGTCTC |
| pos11_Left_HA_FWD | GCTCTGGCTCCGTAGAAGTTG |
| pos11_Right_HA_REV | TGGGCACGTGATGGGAGATA |
| Pos11_noInsert_FWD | CTCAACATGCCCCTCTCCTG |
| Pos11_noInsert_REV | TTTGAAGCTGAGGAGCGACA |
| pos47_Left_HA_FWD | TTGGAATTCTCAGTTCCATCACA |
| pos47_Right_HA_REV | ACTGAGGGGAGGCTTTTAACT |
| Pos47_noInsert_FWD | TCCCCCAATTCTGCAGGTTC |
| Pos47_noInsert_REV | GGCCAGTCTCAGTGGTTCAT |
| pos100_Left_HA_FWD | CTAAGAGAAACAAACGCCAAC |
| pos100_Right_HA_REV | AAGATGAATTGAAAGGAGGTC |
| Pos100_noInsert_FWD | CAGCATCACTGGCCTAACCT |
| Pos100_noInsert_REV | CTCTTCTGCCACCAGGCTAT |
| pos407_Left_HA_FWD | TCATACCAACAGGATCACAAAACA |
| pos407_Right_HA_REV | TGACTGATTCAACAGGGTGCTTT |
| Pos407_noInsert_FWD | TCATACCAACAGGATCACAAAACA |
| Pos407_noInsert_REV | TGACTGATTCAACAGGGTGCTTT |
| posdsRed_Left_HA_FWD | TGGCTGAAATAGGGCAGCAT |
| posdsRed_Right_HA_REV | CGTCACTTACATTTTCCACTGC |
| small_domain_FWD | CCCCTAAGCTATCAGGTTGATTGA |
| small_domain_REV | TCCTTACTGTAGCCTGTGGA |

*ChIP-qPCR*

| | |
|---|---|
| SMC1_pos_control_FWD | CTGAAGATCCCCTGTGCGACC |
| SMC1_pos_control_FWD | ACTACCCAAGGGGATCGAAGC |
| Left_boundary_FWD | TGAAACTTGTGGACATGCCTTATC |
| Left_boundary_REV | AAGAACTGCATTGTGGGATGGAA |
| 3xhCTCF_FWD | TTCAGTCCTTTAGCGCCACC |
| 3xhCTCF_REV | GTCAAGCCACCTGCAAACTG |
| 3xmCTCF_FWD | CTCTTCTGCTCCACCTGCAA |
| 3xmCTCF_REV | AGCCTTAGAGAGGGGTCCAG |
| HBG1_promoter_FWD | TTCAGGGTCAGCTTGCCGTA |
| HBG1_promoter_REV | ACACTCGCTTCTGGAACGTC |

*4C primers Ectopic viewpoints*

| | |
|---|---|
| GFP_FWD | GGGGCACAAGCTGGAGTA |
| GFP_REV | GCTCCTGGACGTAGCCTTC |
| uLCR_FWD | TTTAATATGCTTTAAGTTCTGGGGTAC |

| | |
|---|---|
| uLCR_REV | CTGACCCCGTATGTGAGCA |
| 3xhCTCF_FWD | GGAATCTCGCTCTGATCGTC |
| 3xhCTCF_REV | TGCAGGTGGCTTGACTG |
| 3xmCTCF_FWD | ATTCTCTGCTAGTGTGGGCATC |
| 3xmCTCF_REV | GACCCCTCTCTAAGGCTGACA |

*4C primers Endogenous viewpoints*

| | |
|---|---|
| Left_boundary_FWD | ATCAAAAATGAGTGAAAGGT |
| Left_boundary_REV | CTAGTTGCTTCGTGTGTTCA |
| Pos0_FWD | TTGGCGGCTGCAAGATAC |
| Pos0_REV | AGACCCCCAAACTCCGATT |
| Pos11_FWD | GGAAACCTCAGGACTAGGCAT |
| Pos11_REV | CCCTGATGGTATCGCTGAAT |
| Pos47_FWD | AAGGAGAACAAGGCCTTCAGTA |
| Pos47_REV | TCCTTCCAACCAGCACTCATAG |
| Pos100_FWD | ACAGGGTCTCACTCTGTGGAG |
| Pos100_REV | TTACATTTATATGTGCACAGCAAGTC |
| Pos407_FWD | GGGGTTTCCAGGCAAGTA |
| Pos407_REV | TTGCCATTTCCACCAAGGTC |

**Table S2. oligos for sgRNA cloning**

*For targeting regulatory elements*

| | |
|---|---|
| pos-50_upper | caccgTCAAAGCTGTGCGTCAAGTT |
| pos-50_lower | aaacAACTTGACGCACAGCTTTGAc |
| 3'GFP_cut#1_upper | caccgTACCTGTGGATGTATTTCAA |
| 3'GFP_cut#1_lower | aaacTTGAAATACATCCACAGGTAg |
| GFP_middle_upper | aaacCGGCGCGGGTCTTGTAGTTGc |
| GFP_middle_Lower | caccgAGCACTGCACGCCGTAGGTC |
| 3'GFP_cut#2_upper | caccGTCAATTGGAGGTGTACCTG |
| 3'GFP_cut#2_lower | aaacCAGGTACACCTCCAATTGAC |
| pos0_upper | caccgCACCGTCACCGCCGACGTCG |
| pos0_lower | aaacCGACGTCGGCGGTGACGGTGc |
| pos11_upper | caccgTTCTGAACCAAAGGGGTGCC |
| pos11_lower | aaacGGCACCCCTTTGGTTCAGAAc |
| pos47_upper | caccGGTAGCTACTTCTATAGAGC |
| pos47_lower | aaacGCTCTATAGAAGTAGCTACC |
| pos100_upper | caccGAAACCTCACCGTTTTCAGG |
| pos100_lower | aaacCCTGAAAACGGTGAGGTTTC |
| pos407_upper | caccGGTAGACAACAGGGTACTTG |
| pos407_lower | aaacCAAGTACCCTGTTGTCTACC |
| posdsRed_upper | caccgCCCCTGATTTGGTGCATGGC |
| posdsRed_lower | aaacGCCATGCACCAAATCAGGGGc |
| E407_downstream_upper | caccgATTCATTGGCCTCTCCGTTT |
| E407_downstream_lower | aaacAAACGGAGAGGCCAATGAATc |
| dsRed_upstream_upper | caccgTGACTTTGGGGGTCTAGAAT |
| dsRed_upstream_lower | aaacATTCTAGACCCCCAAAGTCAc |
| Right_boundary_upper | caccgAATTTTCACAAAATTCGGAC |
| Right_boundary_lower | aaacGTCCGAATTTTGTGAAAATTc |

*For KRAB silencing*

| | |
|---|---|
| sg_non-targeting_Control | ttgGTGCACCCGGCTAGGACCGGgtttaagagc |
| sg_non-targeting_Control | ttagctcttaaacCCGGTCCTAGCCGGGTGCACcaacaag |
| sg_SMC1A_upper | ttgGGCCGTACGCCCGAGAACTGgtttaagagc |
| sg_SMC1A_lower | ttagctcttaaacCAGTTCTCGGGCGTACGGCCcaacaag |
| sg_SMC3_upper | ttgGGGAGCGAGCGGCGCTTTGGgtttaagagc |
| sg_SMC3_lower | ttagctcttaaacCCAAAGCGCCGCTCGCTCCCcaacaag |
| sg_RAD21_upper | ttgGAAGGAGGCGCCGGCTGTGGgtttaagagc |
| sg_RAD21_lower | ttagctcttaaacCCACAGCCGGCGCCTCCTTCcaacaag |
| sg_MED21_upper | ttgGGTTTGCTGCGGTAGGAACAgtttaagagc |
| sg_MED21_lower | ttagctcttaaacTGTTCCTACCGCAGCAAACCcaacaag |
| sg_GATA1_upper | ttgGTGAGCTTGCCACATCCCCAgtttaagagc |
| sg_GATA1_lower | ttagctcttaaacTGGGGATGTGGCAAGCTCACcaacaag |
